# Selective pharmaceutical inhibition of PARP14 mitigates allergen-induced IgE and mucus overproduction in a mouse model of pulmonary allergic response

**DOI:** 10.1101/2021.06.05.447208

**Authors:** Alex M. Eddie, Kevin Chen, Laurie B. Schenkel, Kerren K. Swinger, Jennifer R. Molina, Kaiko Kunii, Ariel L. Raybuck, Heike Keilhack, Mario Niepel, R. Stokes Peebles, Mark R. Boothby, Sung Hoon Cho

## Abstract

The type 2 cytokines IL-4 and IL-13, which share use of an IL-4 receptor alpha chain and its nuclear induction of the transcription factor STAT6, are crucial in elicitation and maintenance of allergic conditions that include asthma. Prior work has shown a physical and functional association of STAT6 with PARP14, an ADP-ribosyl monotransferase. Moreover, elimination of all PARP14 expression by gene targeting led to altered recall antibody responses and attenuation of ovalbumin-specific allergic lung inflammation with no apparent health issues for mice lacking this protein. However, an unanswered question is whether or not inhibition of the catalytic function has any biological consequence since PARP14 has multiple functional domains apart from the portion that catalyzes ADP-ribosylation. As reported separately, iterative structural analyses and medicinal chemistry fostered the generation of a compound, RBN2759, that is highly selective in its inhibition of PARP14 with negligible impact on other members of the PARP gene family. We show here that administration of this compound to mice previously sensitized to the allergen *Alternaria alternata* achieved biochemically active levels and altered physiological responses to the antigen. These results show for the first time that in vivo administration of a specific inhibitor of the ADP-ribosyltransferase activity encoded by PARP14 is sufficient to alter biological responses. Specifically, the orally absorbable pharmaceutical compound decreased allergen-induced mucus, blunted the induced increases in circulating IgE, and prevented suppression of IgG2a. We conclude that the catalytic activity can contribute to pathogenesis in allergic processes and propose that other biological endpoints that depend on ADP-ribosylation by PARP14 can be targeted using selective inhibition.

## Introduction

Asthma, including allergen-induced forms of reversible obstructive airways disease, is a highly prevalent condition (1–4). For example, approximately 25 million people in the US (5.5 million children and 19.2 million adults) exhibited evidence of this disease in a self-report survey (1, 4). The standard of care in asthma – which includes inhaled corticosteroids that act on airway inflammation through glucocorticoid nuclear receptors – commonly does not suffice to prevent severe asthma attacks upon re-exposure to environmental allergens (5–7). Although the rates of complete resistance to corticosteroids are low, even partial resistance can undermine long-term health for asthmatic patients, making identification of additional treatments desirable. In addition, strict adherence to regimens of inhaled corticosteroids has remained problematic: a meta-analysis of research from 1985 to 2012 showed no significant improvements in overall clinical success (5–9). Thus, despite a number of therapeutic advances and ongoing introduction of new agents to complement inhaled glucocorticoids, a large burden of morbidity remains in which manifestations of the disease are inadequately controlled, with over 3500 deaths caused by asthma were reported in 2017 (4).

Much remains to be determined about the origins of allergic asthma and pathophysiology of established disease. Although each state can exist independent of the other, there is substantial overlap of asthma with prior atopy (10–13). Atopic or T2-endotype asthma patients typically exhibit an increase in circulating IgE, which points to a contribution of the T helper subset 2 (Th2) of CD4^+^ T cells that secrete type 2 cytokines such as IL-4, IL-5, and IL-13 inasmuch as IL-4 receptor stimulation is crucial for generation of IgE (13–16). Within human asthma and animal models of the disease pathogenesis, eosinophilic (Th2-like) and/or neutrophilic (Th17-like along with Th2) patterns of allergic lung inflammation (ALI) have been identified in patients and mouse models (17–20). Of these, the eosinophilic-pattern disease has seen therapeutic advances with parenteral administration of biologic drugs (e.g., anti-IL-5) whereas the neutrophilic pattern is more associated with severe or glucocorticoid-resistant disease (21, 22) and has seen less progress. Among unmet needs, progress on orally absorbable agents that could mitigate allergic disease or asthma remains important.

During allergic responses, IL-4 and IL-13 production increase due to expansion of the population of allergen-specific Th2 lymphocytes as well as an expansion of group 2 innate lymphoid cells (ILC2). These pleiotropic cytokines act via IL-4Rα receptor complexes on cell targets that include B lymphocytes, alternatively activated macrophages, and airway cells in the lung (23–30). The transcription factor STAT6, induced by IL-4/IL-13 binding to IL-4Rα, plays a critical role in T cell differentiation to functional subsets, chemokine secretion by lung epithelial cells, and in promoting pathological levels of mucus production (28, 30). Moreover, the choice of immunoglobulin (Ig) heavy chain constant region class during class switching in B cells is directed by IL-4, which is essential for generation of IgE, can promote IgG1, and suppresses selection of the Cγ2a isotype for IgG2a production (31). The switch to IgE is directed by induction of the promoter for a non-coding germ line transcript (GLT) termed Iε, which is inducible by IL-4 and initiated by binding of STAT6 to the Iɛ promoter (32, 33). While normally present at low levels, IgE is substantially increased by atopy and in allergic inflammation, during which it functions in the pathophysiology by cross-linking FcɛRI on mast cells and basophils (34).

A screen for transcriptional cofactors that modulate the function of STAT6 in these diverse processes identified the poly (ADP-ribose) polymerase PARP14 (alternatively termed ARTD8) as an interaction partner that binds to the activation domain of STAT6 but not the IFN-γ-induced transcription factor STAT1 (35). Analyses of mice lacking detectable expression of any PARP14 after gene inactivation provided evidence that allergic lung inflammation provoked by the model antigen ovalbumin and ovalbumin-specific IgE responses are decreased in the absence of PARP14 (36, 37). Moreover, the domain of PARP14 homologous to mammalian PARP1 and to ADP-ribosyltransferases (ARTs) encoded by bacterial exotoxins is enzymatically active (38). Unlike the members of the PARP gene family that function as processive polymerases, PARP14 is thought to transfer only one or a few ADP-ribose moieties onto protein acceptor sites (39). However, in addition to the ADP-ribosyltransferase catalytic domain, PARP14 contains several other functional modules (35, 40, 41). As such, whether or not there are functional consequences of the ADP-ribosylation executed in vivo by this protein, or if so what are they, remain unanswered questions.

One approach to allow identification of physiological functions of PARP14 in the intact animal would involve administration of a highly selective inhibitor of its ADP-ribosylation activity. This approach was not realized for many years due to challenges inherent in generating such a compound (42–44). The post-translational modification of proteins by ADP-ribosylation involves the transfer of ADP-ribose (ADPr) from NAD^+^ to an amino acid acceptor in a target protein. Mammals express extracellular as well as intracellular ADP-ribosyltransferases (ARTs), and also encode a variety of enzymatic activities that remove ADPr from modified proteins (45–47). Intracellular ARTs in mammals include those that can form branching polymers by iterative additions of ADPr to ADPr, i.e., polymerases or poly-PARPs (e.g., PARP1 and 2), and those that do not polymerize, i.e., mono-PARPs or –ARTs such as PARP14 (45–47). Among the 17 intracellular PARP family proteins, the NAD-binding pockets are structurally similar. Attempts to use existing PARP inhibitors as surrogates suffer from the drawbacks that their IC50 for PARP14 requires the use of concentrations that both are more inhibitory for poly-PARP enzymes such as PARP1 and PARP2 and from cellular toxicities at the concentrations affecting PARP14 (36, 48). Moreover, *Parp1* gene inactivation studies as well as use of PARP1, 2-specific inhibition using olaparib suggest that ovalbumin-induced allergic lung inflammation depends on the polymerase PARP1 (49–53). This confounding variable makes interpretations of results with broad-spectrum inhibitors such as PJ-34, which only weakly inhibits PARP14, problematic because the coverage of PARP1 will exceed that of PARP14. Recently, however, RBN2759 has been identified as an orally absorbable small molecule that selectively inhibits the catalytic activity of PARP14 with workable pharmacological properties (54, 55). Accordingly, we tested if administration of this compound could alter any functional aspects of the immune response to an aeroallergen.

To do so, we selected a mouse model that entails administration of an asthma-relevant aeroallergen. Airway inflammation is elicited with a known aeroallergen, *Alternaria alternata*, in one class of such models. This fungal species elicits an IgE-mediated respiratory disease and is a trigger for flares of asthma and allergic disease in patients (56, 57). Because therapeutic potential of a pharmaceutical in patients hinges on its effect after initial immune experience, we tested how the highly selective PARP14 inhibitor influenced IgE levels, mucus formation, and other features of allergic response elicited by inhaled rechallenge with *Alternaria* when administered after a phase of priming by serial inhalation.

## Materials & Methods

### Mice

Male and female BALB/c-J (Jackson Laboratories) mice were housed in ventilated micro-isolators under specified pathogen-free conditions in a Vanderbilt University Medical Center mouse facility and used at 6-8 wk of age following protocols approved by the Institutional Animal Care and Use Committee.

### Immunization and pharmaceutical treatment

To prime mice for recall allergic response, *Alternaria* extract (lot # 338869, Greer Laboratories, Lenoir, NC) was administered in PBS (5 μg in 50 μL) was administered by intranasal (i.n.) instillation once daily for five days. Two weeks later, mice were re-challenged once daily for three consecutive days with the same dose and route of administration. RBN-012759, hereafter RBN2759 (PARP14 inhibitor, i.e., PARP14i) (55), a compound generated by Ribon Therapeutics, Inc. (Cambridge MA), was delivered in a vehicle of 0.5% methylcellulose, 0.2% Tween 80 dissolved in sterile water. Four days prior to *Alternaria* rechallenge, mice began receiving PARP14 inhibitor (500 mg/kg) by oral gavage twice daily. Peak and terminal trough plasma samples were collected two hours after the penultimate gavage and the day following the final intranasal challenge and gavage, respectively, by using 15% (w/v) potassium EDTA. Frozen portions of these samples were used for Ab assays and pharmacokinetic analysis. Levels of RBN2759 compound in plasma samples at these times were analyzed Charles River Laboratories using liquid chromatography and mass spectrometry. The left mainstem bronchus of each harvested mouse was clamped to isolate the left lobes, followed by bronchoalveolar lavage (BAL) performed on the right lung lobes using sterile PBS (700 μL). The right lung lobes were used to prepare a cell suspension by digestion with collagenase (1.5 mg/mL) and hyaluronidase (1.0 mg/mL) as described (36), followed by flow cytometry and culture. Left upper lobes were placed in formalin overnight then embedded in paraffin. The remaining tissue was snap frozen and stored at −80°C for later isolation of RNA.

### Histology, bronchoalveolar lavage fluid, and cytometry

Serial sections (5 μm thickness, positioning two sections per slide) of formalin-fixed, paraffin-embedded lung tissue were deparaffinized with xylene, and then stained using Periodic Acid Schiff (9162B, Newcomer Supply, Middleton WI). To score airway mucus semi-quantitatively, three fields were selected for each sample in a manner masked as to sample identity. Images of all fields were then scored independently by three individuals masked as to sample identity. A scale quantified as 0-3 (0, no excess mucus; 1, marginal and occasional hyperplasia and mucus; 2, substantial and moderate mucus cell hyperplasia and some airway mucus; 3, severe hypersecretion and airway plugging) was used, and average scores for a given sample were generated from the three individual scores, which most often were perfectly concordant and at most differed among each other by 1. The average score for each subject was then used to calculate a mucus hypersecretion index. Differential counts of bronchoalveolar lavage fluid samples, performed by a reader masked as to sample identity, were analyzed after cytospin deposition on microscope slides and Richard-Allen Three-Step staining [#3300, Thermo-Fisher (58)].

### Measurements of cytokines and Ab

Secreted IL-4 was measured using a matching antibody pair (capture, Tonbo 70-7041-U500; detection, eBio # 13-7042-85) with color generated using streptavidin-HRP (R&D Systems, Dx998) followed by Ultra-TMB reagent (Pierce; Thermo #34028). Both the purified and biotinylated Abs were used at a concentration of 0.5 μg/well. Supernatants were collected from 1.0 × 10^6^ lung cells/mL media restimulated overnight with plate-bound anti-CD3 (adsorbed to wells at 1.0 μg/ml in PBS) and soluble anti-CD28 (1.0 μg/ml) (Tonbo Biosciences), as previously described (36, 59). Dilution points for the ELISAs were selected based on the linear range of standard curves with purified recombinant cytokine. Cytokine levels in restimulated lung suspension also were measured by Th1/Th2/Th17 Cytometric Bead Array (BD 560485). Relative concentrations of circulating antibodies were determined using plasma collected at the time of harvest. High affinity binding plates were coated with purified Ab (0.5 μg/well) directed against Ig H+L (Southern Biotech) for IgG classes or IgE (BD Pharmingen, #553413) or with *Alternaria* extract (0.1 μg/well). Quality control testing indicated that only anti-*Alternaria* IgG1 yielded reliably specific signal.

### Gene Expression

Total RNA was extracted from frozen lung tissue using TRIZOL (Invitrogen) and a Micro-BeadBeater 96 (Biospec). RNA concentration and purity were measured using a NanoDrop. cDNA were synthesized from RNA (4 μg) using AMV Reverse Transcriptase (Promega)., as described (36). Gene expression was quantified using PowerUp SYBR green master mix (Qiagen, Valencia, CA) via quantitative real-time (qRT)-PCR. Mucus-producing hyperplasia was estimated using *Muc5ac* primers with gene expression normalized to β-actin.

#### Immunoblotting for detection of ADP-ribosylation

Protein was extracted from splenocyte suspensions using modified 10 mM Tris pH7.4, 150 mM NaCl, 2 mM EDTA, 1% NP-40, 1% sodium deoxycholate and 0.1% SDS, supplemented fresh with 0.1 M Master Protease Inhibitor (P8340, Sigma-Aldrich). Mutants of full-length PARP14 that lack ADP-ribosyltransferase catalytic activity as well as a construct encoding the wild-type protein were described previously (36). Lysates of ΦNX cells transfected with WT of pcDNA3-FLAG-PARP14 were either analyzed directly or used for immune precipitations with monoclonal anti-FLAG (M2) (Sigma-Aldrich). The resulting immune complexes, collected using protein G beads (Santa Cruz Biotechnology, Santa Cruz, CA), were rinsed, and eluted proteins were analyzed by immunoblotting. Filters were blocked with 5% milk in 0.05% Tween20 and incubated (2 hr at 20 C) with primary antibodies directed against the FLAG epitope tag, mouse actin, mouse PARP14, and PAR/MAR (Cell Signaling Technologies #83732S) and rinsed. After incubation with secondary antibodies (IRD800 or IRD680) and rinsing, indirect immune fluorescent bands were visualized and quantitated using an Odyssey imaging system (Li-Cor, Lincoln NE) as described (59).

### Cell cultures

B cells were purified (90–95%) by negative selection with single-cell splenocyte suspensions by using biotinylated anti-Thy1.2 mAb followed by streptavidin-conjugated microbeads (BD Pharmingen). In brief, 0.75 x10^6^ cells/mL were cultured in 3 mL of RPMI1640 medium (Gibco Cat. #23400-021) in a 6-well cell culture plate (Peak Serum Inc. Cat. #TR5000), supplemented with recombinant mouse BAFF (10 ng/mL). B cells were activated and plasma cell differentiation stimulated using anti-CD40 (BD Cat. #553788) (1 μg/mL) or LPS (Sigma L2630) (1 μg/mL), supplemented with IL-4 (Peprotech Cat. #214-14) (10 ng/mL) and IL-5 (Peprotech Cat. #215-15) (10 ng/mL) as indicated. RBN2579, dissolved in 100% DMSO for stock solutions (1 mM; 368 μg/mL). Cultures were treated with either DMSO or PARP14i (1 μM or 0.33 μM RBN2759), added once daily. Cultures were harvested and counted on Day 5; supernatants were collected for ELISA to quantify relative levels of antibody (IgG1, IgG2a, and IgE).

### Statistical Analysis

Graphs and statistics were generated using PRISM software (GraphPad, San Diego, CA). Averages were generated from results in five independent experiments (three with male mice and two with females) with no exclusions of subjects; drop-out was only for spontaneous mortality in the course of the experiment, the rate for which was not affected by treatment with active compound as compared to vehicle. Two-way ANOVAs were performed on the datasets across full dilution curves, followed at specific points on the curves by either a Student’s T-test or Welch’s unpaired T-test, selected according to the variance between the two populations for which the null hypothesis was tested.

## RESULTS

### PARP14-specific inhibitor RBN2759 attenuated ADP-ribosylation in vivo in a mouse model of *Alternaria*-induced allergic responses

A model of allergy induction and adaptive immune recall was used in which mice first were primed by daily inhalations of *Alternaria* antigens (Ag) in a sensitization phase. After 12 d, the subject animals started to receive either vehicle or PARP14 inhibitor (RBN2759) followed after four days by five consecutive daily challenges with inhaled Ag while continuing twice-daily gavages with inhibitor or vehicle (Fig. 1A). To assess the circulating concentrations of drug during treatment, plasma samples were collected at 2 hr after a dose and at trough concentration just prior to morning gavage (14 hr samples). Quantitation of the compound by LC-MS revealed mean concentration of ~ 3.3 μM (~1227 ng/mL) at 2 hr post-dose, albeit with significant variance (range, 497 to 1680 ng/mL) (Fig. 1B). Mean trough concentrations were 237 ng/ml (~0.64 μM; range, <50 to 1030 ng/mL), with over half of the subjects having a concentration ≥ 0.33 μM (Fig. 1B). No specific target of PARP14 has been well established in vivo; moreover, it clear not if the general ADP-ribosylation of any such target is exclusively catalyzed by PARP14 inasmuch as several other ADP-ribosyl monotransferases and polymerases are expressed concurrently. Accordingly, we first validated the inhibitory capacity of RBN2759 by transfecting cells with PARP14 expression vector (36) or a control, culturing in inhibitor or vehicle, and immunoblotting extracted proteins with antibody recognizing mono-as well as poly-ADP-ribose adducts on proteins (MAR and PAR, respectively) (Fig. 1C). This analysis showed that inhibitor treatment substantially reduced the auto-modification of over-expressed PARP14. In light of the observed inhibition, we tested if the treatment of mice with orally delivered RBN2759 decreased the intensity of anti-MAR/PAR bands in splenocytes of subject mice. This analysis revealed multiple bands reproducibly decreased in samples from inhibitor-treated subjects compared to controls (Fig. 1D). PARP14 expression is increased by pro-inflammatory signals such as TNF-α (41). This suggested a biological impact of inhibiting ADP-ribosylation in *Alternaria*-exposed lung tissue should cause reduced PARP14, which was observed (Fig. 1E). We conclude that this regimen of oral dosing of mice with the tool compound RBN2759 achieved active concentrations and had a biochemical impact in vivo, with lower PARP14 levels and less ADP-ribosylation of target proteins in tissues of the treated mice.

**Fig. 1.**
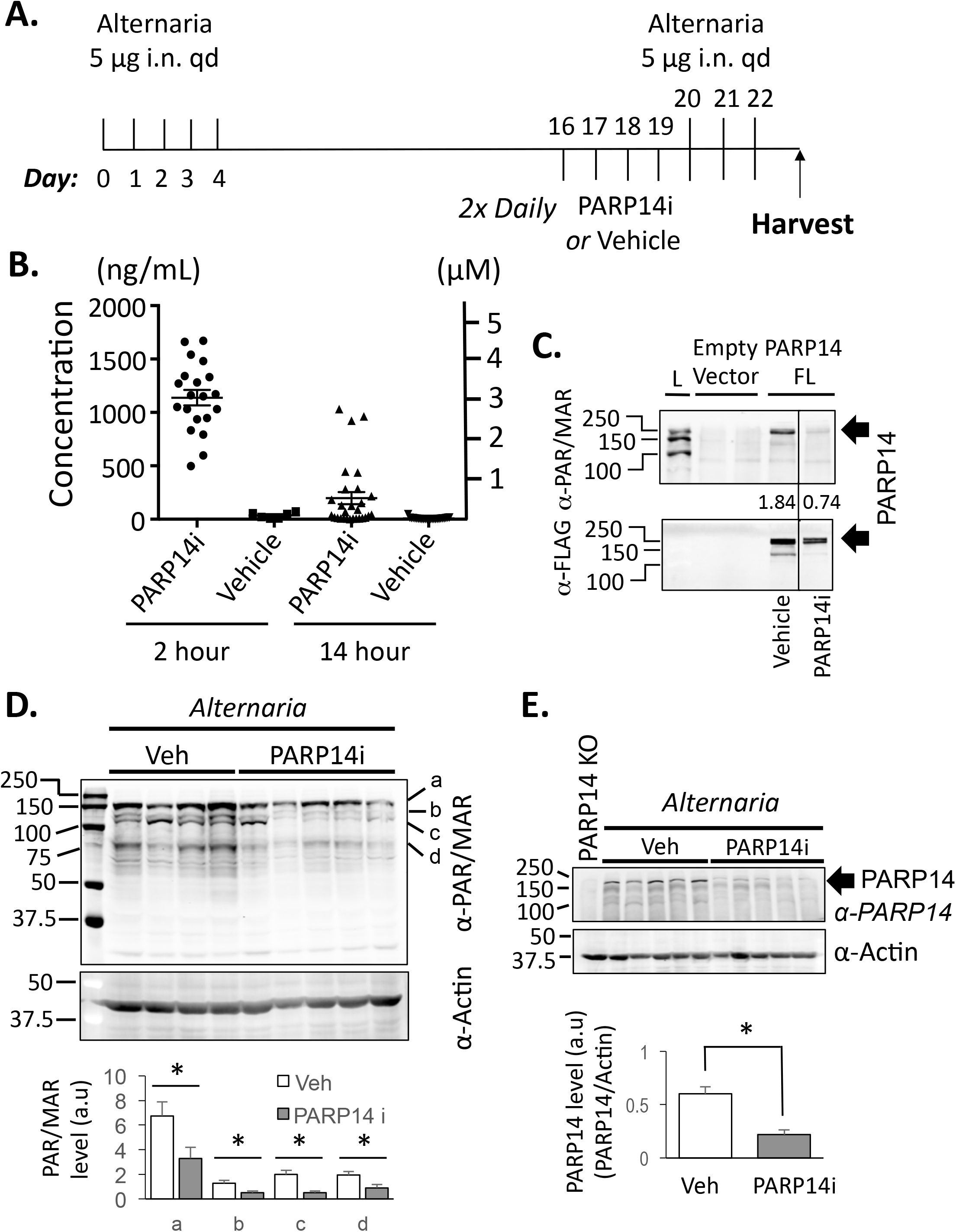
In vivo biochemical effect from an inhibitor of PARP14-mediated ADP-ribosylation in an allergic disease model. (A) Timeline of airway sensitization with *Alternaria* extract (5 μg per dose), treatment with the selective PARP14 inhibitor (PARP14i) RBN2759 (500 mg/kg/dose) or vehicle, and intranasal recall challenges. (B) Concentrations of RBN2579 in terminal plasma from mice administered gavages of vehicle or PARP14i. Shown are results of LC-MS measurements after the final dose before harvest (~14 hour post-*gavage*; n= 28 vs vehicle, n= 25), or after the penultimate dose of PARP14i (2 hour; n= 20), vehicle n= 8). (C) ФNX cells transfected were transfected with pcDNA3 with no insert or FLAG epitope-tagged PARP14 and cultured in vehicle or RBN2759 (1 μM) and extracted proteins were immuno-precipitated with anti-FLAG. Shown are results of immunoblots probed with αPAR/MAR and αFlag (upper, lower images, respectively). Arrow indicates the band of interest for each respective immunoblot. The values shown are αPAR/MAR band intensities normalized to the band intensity of αFLAG. (D) Altered ADP-ribosylation in ex vivo tissue after administration of RBN2759. Female mice were sensitized (8 d) with *Alternaria* extract (3 μg in 50 μL daily) while administered gavages of vehicle or compound (500 mg/kg twice per d). Single-cell suspensions of spleens from mice sensitized daily with *Alternaria* extract while receiving RBN2759 twice daily by gavage were prepared ~15 h after the last gavage and analyzed by immunoblotting with antibodies directed to the indicated ligands after resolution of unfractionated lysates on SDS-PAGE. Letters (a-d) indicate the positions of bands reproducibly decreased with PARP14 inhibition in vivo, quantitation (means ± SEM) of which is shown in bar graphs below the gel image. (E) As in (D) except that an extract of B6-*Parp14* −/− splenocytes was included and lysates were probed with antibody against PARP14, as described (35, 36). (D, E) The likelihood of the null hypothesis for indicated differences (*) was p<0.05.

### Treatment with RBN2759 mitigated *Alternaria*-induced mucus accumulation

Mucus plugging and hyper-production are major factors in the airflow obstruction in human asthma (60, 61) so we had selected this parameter as the primary end-point in analyses of the allergen-rechallenged mice. In addition, because of sex differences in responses of mice to allergens such as *Alternaria alternata*, we separately analyzed male versus female mice. The histologic results showed that, as expected, allergen sensitization was essential for positive scores (i.e., when comparing *Alternaria*-challenged mice that received vehicle by gavage) (Fig. 2A, B). When compared to vehicle controls, pre-treatment with the inhibitor specific for PARP14 catalysis caused a reduced amount of PAS-positive cells in the airways as well as mitigating mucus in the airways of sensitized and challenged mice (Fig. 2A, representative samples; Fig. 2B, aggregate data for all mice). Male and female mice exhibit quantitatively different responses to *Alternaria*. Although RBN2759 reduced mucus in both males and females, and in a few cases no hyper-secretion was detected, the median effect did not reduce scores to the level of non-sensitized controls. Similarly, when *Muc5a* mRNA levels that encode a protein component of mucus were measured by qPCR with cDNA prepared from the RNA of one lung lobe, these were lower in *Alternaria*-challenged mice treated with active compound when compared to vehicle-treated controls. We conclude that the mucus over-production after challenge with *Alternaria* antigens was reduced by selective in vivo inhibition of the capacity of PARP14 to execute ADP-ribosylation.

**Fig. 2.**
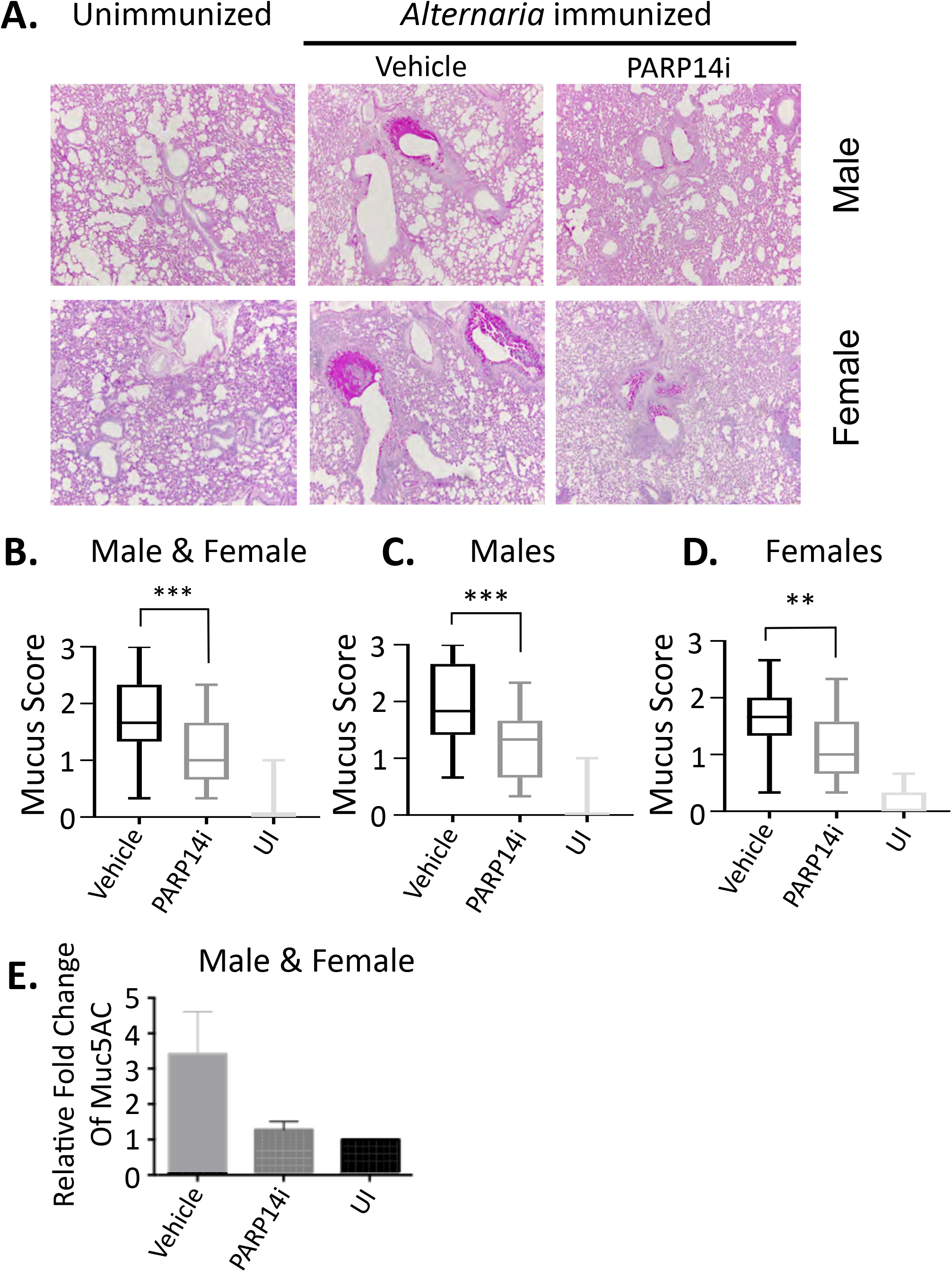
Selective PARP14 inhibitor protects against allergen-induced mucus hyper-production. (A-C) Mice were sensitized and challenged with *Alternaria* extract, and treated with RBN012759 or vehicle, as in Fig. 1A. (A) Representative photomicrographs of individual PAS-stained sections from male and female mice in the indicated treatment groups. (10x Magnification) (B) Selective PARP14 inhibition reduces mucus in *Alternaria*-sensitized mice. Mucus scores were developed from PAS-stained lung sections by analysts blinded to sample identity as detailed in the Methods. Shown are separate panels with mean (±SEM) scores for all subjects, males, and females, as indicated. The probability of the null hypothesis being correct was less than 0.01 (**) or 0.001 (***), as indicated. In each case, scores in allergen-sensitized, rechallenged mice were statistically significant in their difference from non-immune (UI) cagemate controls mice. (C) Expression of *Muc5ac* mRNA in lungs of vehicle- and PARP14i-treated *Alternaria*-allergic mice after antigen re-challenge was analyzed by qRT-PCR of RNA extracted from lungs, normalized to the mean *Muc5ac* mRNA in the lungs of unimmunized mice.

### Effect of RBN2759 on *Alternaria*-induced immune cell profiles uncoupled from mucus reduction

In analyses of broncho-alveolar lavage (BAL) of patients with active disease, atopic asthmatics commonly have increased eosinophils in their airways (62, 63); a similar effect is typical in mouse models of allergic lung inflammation. Accordingly, we analyzed differential counts of the cells recovered by unilateral BAL after clamping one mainstem bronchus. As expected, the fractions of recovered cells that were eosinophils were substantially greater in *Alternaria*-sensitized and rechallenged mice (Fig. 3). The type 2 inflammatory cytokines IL-4 and IL-13 are major drivers of mucus over-production, and eosinophil recruitment and survival are promoted by IL-4 and the type 2 cytokine IL-5, respectively (14–16, 30, 64). Surprisingly, however, no effect of the PARP14 inhibitor was observed for the BAL samples from the same mice as clearly had reduced mucus (Fig. 3). Some evidence suggests that inhibition of the drivers of type 2 inflammation may increase Th17 responses or the neutrophil recruitment that is enhanced by IL-17 (65). However, RBN2759 treatment at this dose did not change frequencies of neutrophils in the BAL fluids recovered from *Alternaria*-challenged mice. Such mouse models often also have increased immune cell infiltration in the parenchyma. Numbers of cells in lobar dispersions of lungs from mice in these experiments showed no substantial impact of PARP14 inhibitor on the increases elicited by allergic sensitization followed by inhaled rechallenge (Table 1). Although a trend toward modest reduction was observed for the numbers of B cells and neutrophils (PMN) with a modest reduction in CD4^+^ T cells in females, flow cytometric characterizations did not reveal statistically significant changes in major cell types or subsets (Table 1). These changes suggest that inhibition of PARP14 at the recall stage of allergic lung inflammation in vivo may have a modest effect on recruitment of several types of immune cell, but we conclude that the magnitude of decrease in mucus over-production was largely uncoupled from global indices of inflammatory cell recruitment.

**Table 1.**

| % | CD4<sup>+</sup> | CD8<sup>+</sup> | B cell | Macrophage | Eos (Gr1<sup>++</sup>) | PMN (Gr1<sup>+</sup>) |
| --- | --- | --- | --- | --- | --- | --- |
| UI-Veh. | 27.0 ± 0.3 | 12.4 ± 0.7 | 22.3 ± 0.2 | 15.1 ± 8.6 | 3.1 ± 1.6 | 26.4 ± 16.6 |
| UI-PARP14i | 32.0 ± 3.0 | 10.6 ± 0.7 | 25.8 ± 4.1 | 14.22 ± 10.9 | 9.0 ± 3.2 | 19.9 ± 13.3 |
| Veh. | 26.4 ± 1.5 | 9.8 ± 0.4 | 28.8 ± 1.7 | 24.5 ± 3.6 | 2.9 ± 0.4 | 37.9 ± 4.1 |
| PARP14i | 25.5 ± 2.0 | 9.7 ± 0.8 | 25.6 ± 2.3 | 22.1 ± 3.8 | 5.8 ± 2.9 | 32.3 ± 4.7 |

|  | CD4<sup>+</sup> | CD8<sup>+</sup> | B cell | Macrophage | Eos (Gr1<sup>++</sup>) | PMN (Gr1<sup>+</sup>) |
| --- | --- | --- | --- | --- | --- | --- |
| Unimm-Veh. | 2.2 ± 0.4 | 1.0 ± 0.2 | 1.8 ± 0.3 | 1.6 ± 1.1 | 0.3 ± 0.1 | 2.5 ± 1.9 |
| Unimm-PARP14i | 3.0 ± 0.05 | 1.0 ± 0.1 | 2.5 ± 0.6 | 1.4 ± 1.1 | 0.8 ± 0.2 | 2.0 ± 1.4 |
| Veh. | 3.2 ± 0.2 | 1.2 ± 0.1 | 3.8 ± 0.5 | 3.5 ± 0.8 | 0.4 ± 0.06 | 4.7 ± 0.4 |
| PARP14i | 3.1 ± 0.2 | 1.2 ± 0.1 | 3.4 ± 0.6 | 3.1 ± 3.8 | 0.6 ± 0.3 | 4.0 ± 0.6 |

**Females**
| % | CD4<sup>+</sup> | CD8<sup>+</sup> | B cell | Macrophage | Eos (Gr1<sup>++</sup>) | PMN (Gr1<sup>+</sup>) |
| --- | --- | --- | --- | --- | --- | --- |
| Unimm-Veh. | 43.0 ± 4.3 | 16.7 ± 0.5 | 15.6 ± 3.6 | 2.9 ± 1.5 | 1.9 ± 1.4 | 4.6 ± 0.8 |
| Unimm-PARP14i | 39.5 ± 1.8 | 14.9 ± 2.4 | 20.4 ± 1.0 | 2.5 ± 1.3 | 1.2 ± 0.2 | 7.1 ± 0.7 |
| Veh. | 28.3 ± 1.3 | 8.2 ± 0.7 | 30.3 ± 3.8 | 9.1 ± 1.1 | 2.2 ± 0.3 | 29.6 ± 1.1 |
| PARP14i | 27.8 ± 1.5 | 8.4 ± 0.7 | 30.3 ± 2.8 | 9.0 ± 1.2 | 2.5 ± 0.2 | 31.1 ± 1.8 |

|  | CD4<sup>+</sup> | CD8<sup>+</sup> | B cell | Macrophage | Eos (Gr1<sup>++</sup>) | PMN (Gr1<sup>+</sup>) |
| --- | --- | --- | --- | --- | --- | --- |
| UI-Veh. | 2.2 ± 1.6 | 0.9 ± 0.7 | 1.0 ± 0.9 | 0.2 ± 0.2 | 0.04 ± 0.01 | 0.2 ± 0.1 |
| UI-PARP14i | 2.1 ± 1.2 | 0.7 ± 0.3 | 1.2 ± 0.7 | 0.2 ± 0.2 | 0.06 ± 0.03 | 0.4 ± 0.3 |
| Veh. | 4.8 ± 1.4 | 1.3 ± 0.4 | 6.9 ± 2.7 | 1.9 ± 0.7 | 0.5 ± 0.2 | 5.8 ± 1.9 |
| PARP14i | 4.0 ± 1.0 | 1.2 ± 0.3 | 5.4 ± 1.7 | 1.7 ± 0.5 | 0.4 ± 0.1 | 5.1 ± 1.4 |

**Fig. 3.**
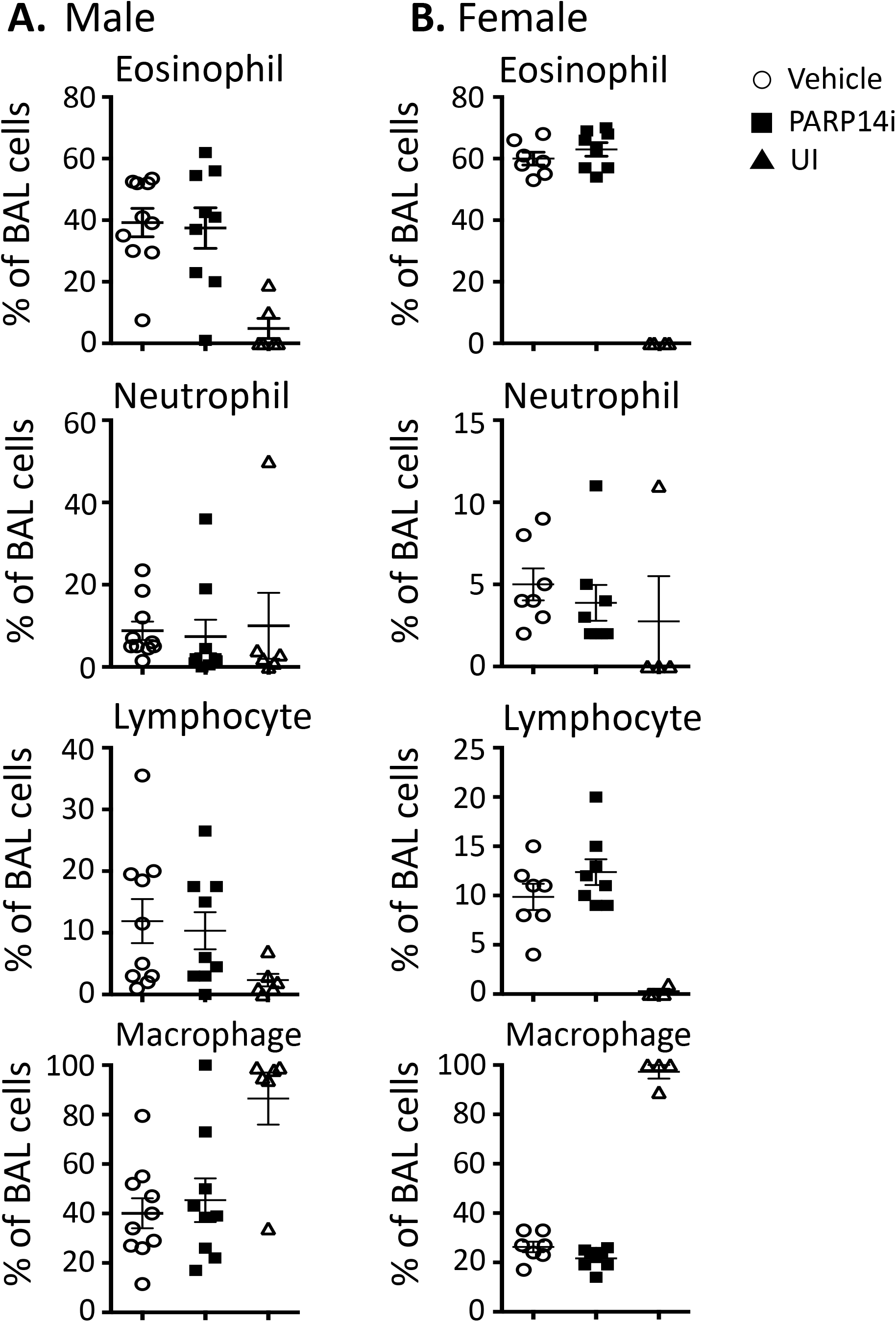
Lack of reduction in inflammatory cells recruited to airspaces. Differential counts were performed on cells spun onto microscope slides from aliquots of recovered BAL fluid after staining. Data are shown separately for males (A) and females (B). Shown are the mean (±SEM) frequencies of the indicated inflammatory cells in the BAL fluid from unimmunized (UI) and *Alternaria-*immunized mice treated with vehicle or PARP14i prior to re-challenge.

### Reduced IL-4 production after TCR stimulation of pulmonary infiltrate

One potential explanation for capacity of a PARP14-selective inhibitor to reduce *Alternaria*-induced mucus accumulation would be that although the extent of infiltration was similar, the amount of type 2 cytokine release would be diminished. To test this possibility, equal numbers of cells in the lung suspensions were stimulated via the TCR to gauge T cell-specific effects. Activation by a combination of phorbol ester (PMA) and calcium ionophore was also performed since these stimuli can also elicit cytokines from mast cells, basophils, and ILC as potential sources of IL-4 or −13. The results show that IL-4 production was dramatically increased by allergic sensitization, and this effect was blunted by the treatment with PARP14 inhibitor (Fig. 4A). Surprisingly, however, this reduction was only observed with male subjects (Fig. 4B, C) and was not a function of the frequency of Th2 cells as judged by flow cytometry after staining of intracellular IL-4. Consistent with the IL-4 data, secreted type 2 cytokines IL-5 and IL-13 also were reduced after re-stimulation of lung cells from male but not female mice (Fig. 4D, E). There is evidence that PARP14, by an unknown mechanism, can promote Th17 differentiation of naïve T cells (36). IL-17 production by the same cell suspensions was also reduced with lung cells from male subjects treated with PARP14 inhibitor after either T cell-specific or broadly acting stimuli (Fig. 4F & G, respectively). Nonetheless, IL-17 production after stimulation of lung cells from RBN2759-treated females was indistinguishable from the controls (not shown). Moreover, production of the T cell-specific cytokine IL-2 and the pro-inflammatory cytokine TNF-α were each lower in the inhibitor-treated males and undetectable in the absence of *Alternaria* sensitization (Fig. 4F & G). These were cytokine-specific effects and not a cellular toxicity in that cytokines such as IL-10 were similar in the samples from treated, vehicle control, and non-sensitized mice (Fig. 4F & G). We conclude that despite the lack of effect on overall numbers of inflammatory cells in the lungs or BAL fluid of inhibitor-treated *Alternaria*-challenged mice, treatment of males with the drug prior to and during re-challenge attenuates type 2 cytokines (IL-4, −5, and −13) as reducing the capacity to produce IL-2, TNF-α, and IL-17.

**Fig. 4.**
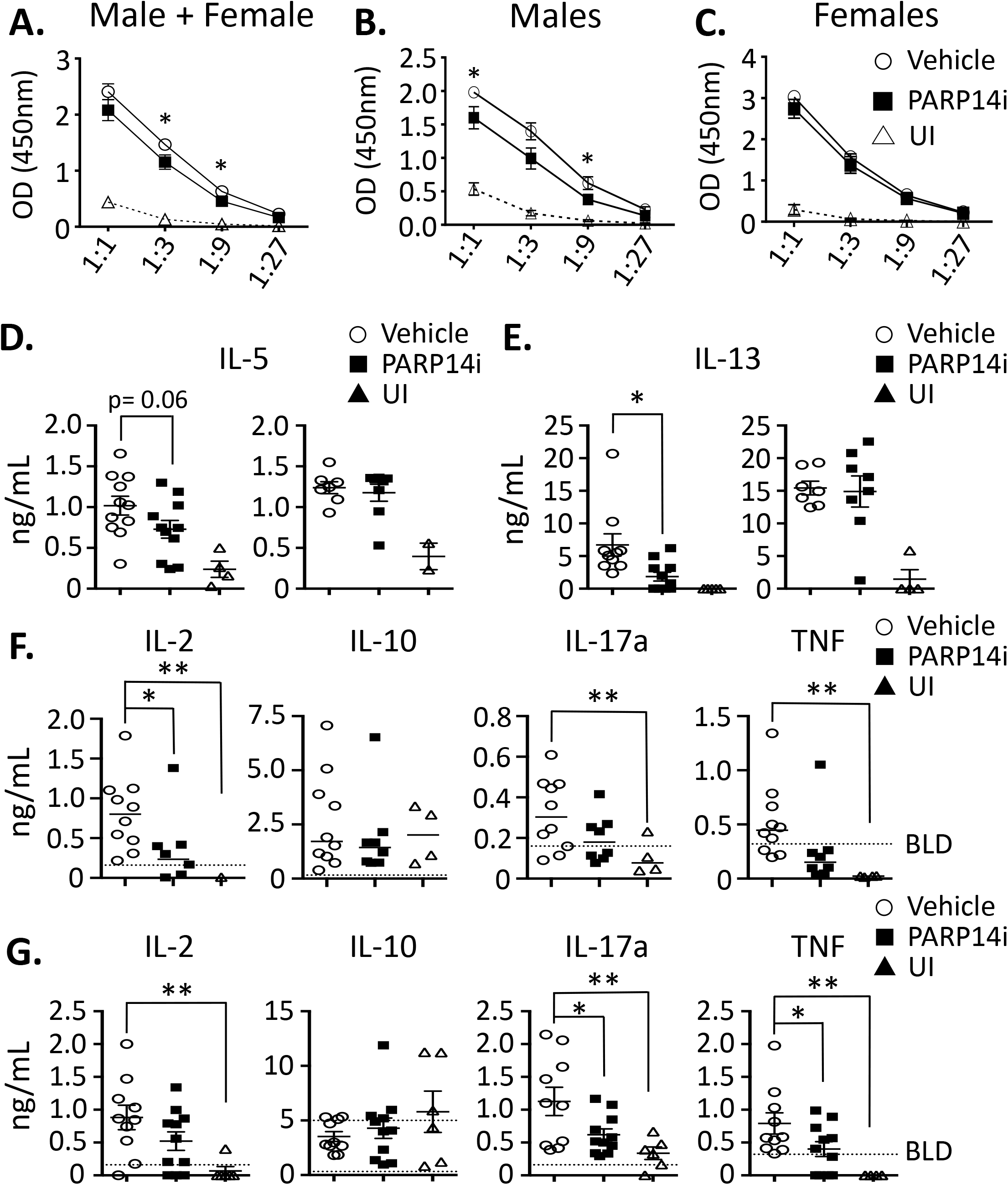
Selective in vivo inhibition of PARP14-catalyzed ADP-ribosylation reduces IL-4 and other cytokine production by pulmonary cells ex vivo. (A-C) Shown are results (means ±SEM) of ELISA to measure IL-4 in culture supernatants after re-stimulation of lung suspensions via the TCR, averaging data for males and females combined (A), males (B), or females (C). (*), the probability of the null hypothesis applying was less than 0.05 at the indicated individual points, and <0.01 by ANOVA across the curves in (A) and (C). (D, E) Measurements on the supernatants assayed by ELISA in (A), but with single-point values interpolated to standard curves for measurement of IL-5 (D) and IL-13 (E) in supernatants from male (left panel of pair) and female samples. Shown are individual values as indicated, with horizontal lines marking mean values. (F) Additional cytokines measured in the same supernatants for male subjects, displayed as in (D). (G) As in (F) except that the supernatants were from lung suspensions that were re-stimulated with PMA and ionomycin. Only male subjects’ data are shown as no meaningful difference between PARP14i and control mice was observed with samples from females. *, p<0.05; **, p<0.01; *** p<0.001 as results of testing null hypothesis.

### Inhibition of allergy-associated increase in IgE

The induction of IgE is one of the first responses to IL-4 to be identified and increased total IgE levels in humans are a hallmark of atopy. In parallel with its promotion of isotype switching to IgH Cγ (32) and Cγ1, this cytokine inhibits switching to Cγ2a/c and levels of IgG2a/c (31). Accordingly, we analyzed the relative concentrations of antibodies of these isotypes in *Alternaria*-challenged mice and non-immunized controls. IgE levels in sera of both male and female mice increased dramatically with allergic sensitization (Fig. 5A-C). This effect was substantially reduced in female as well as male mice pre-treated with RBN2759 when compared to vehicle controls (Fig. 5A-C). A reciprocal effect was observed for IgG2a, whose circulating levels were reduced by allergic sensitization in vehicle-treated mice, whereas the mice subjected to PARP14 inhibition exhibited levels of IgG2a indistinguishable from non-sensitized controls (Fig. 5D). Unlike IgE, the IgG1 isotype is robustly induced even with short-term primary exposure to antigen, and circulating IgG has a far longer half-life than IgE due to selective FcRn-mediated re-uptake of IgG via FcRn. Moreover, although promoted by IL-4, IgG1 induction can be IL-4-independent. Of note, IgG1 levels in sera of inhibitor-treated mice and vehicle controls did not differ (Fig. 5E). Moreover, an ELISA that allowed reliable detection of *Alternaria*-specific IgG1 detected no effect of RBN2759 on specific antigen (Fig. 5F). We conclude that specific inhibition of the ADP-ribosyl monotransferase activity in PARP14 impacted the nature of an antibody response in vivo. Thus, RBN2759 interfered with the capacity to generate the STAT6- and IL-4-dependent IgE isotype upon sensitization and challenge of both males and females, whereas long-lived IgG1 was unaffected.

**Fig. 5.**
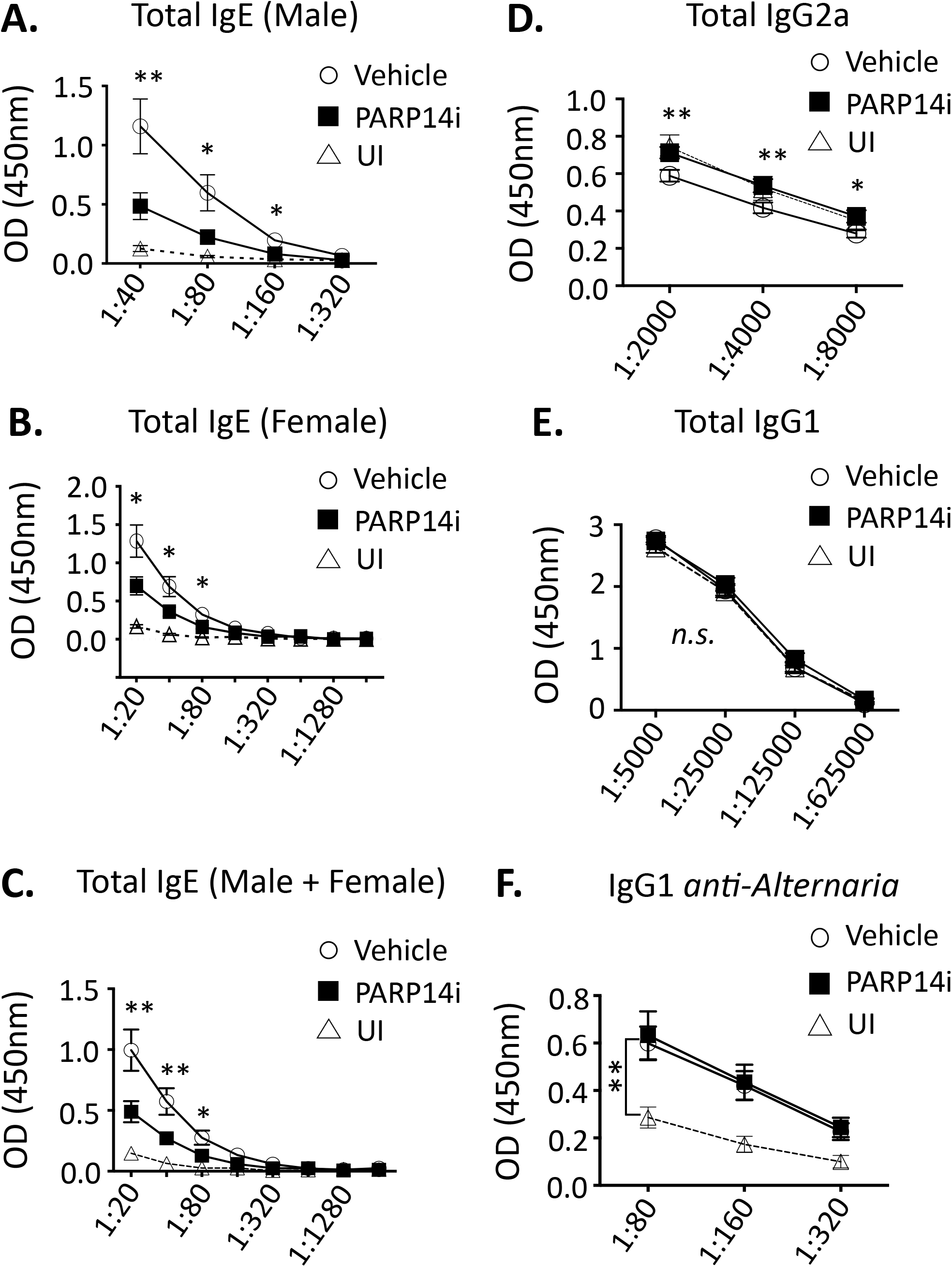
PARP14 inhibitor blunts allergen-induced changes in IgE and IgG2a. (A-F) Terminal plasma samples were collected from unimmunized (n=10), *Alternaria*-immunized PARP14i-treated (n=19), and *Alternaria*-immunized vehicle control (n=17) subjects. Shown are the mean results (±SEM) for total IgE, averaging all samples for males (A), females (B) or all samples (C). (D) Results of ELISA for total IgG2a at dilutions in the linear range of titrations, averaging samples from all subjects (male and female) as in (C). (E) Total IgG1, as in (D). (F) ELISA measuring allergen-specific IgG1 using *Alternaria* extract to capture on plates. Linear range dilutions were selected from standard curve; the vertical tie bar denotes the statistically significant increases in antigen-specific IgG1 over unimmunized control for both inhibitor-treated and control subjects. * p≤0.05; ** p≤0.01; n.s., non-significant.

In addition to reduced IgE responses to ovalbumin in PARP14-deficient mice, in vitro evidence supported both B cell-intrinsic and cell-extrinsic mechanisms (36). To test if the finding of reduced IgE depended on ADP-ribosylation by PARP14, we activated B cells with either LPS or by cross-linking CD40, cultured them in IL-4 with addition either of vehicle of inhibitor, and measured the secreted IgE. RBN2759 reduced IgE production to an extent almost as substantial as the overall effect in males in vivo (Fig. 6A, B), including at 0.33 μM (Fig. 6E), the concentration that was the median trough level in vivo (Fig. 1B). Relative IgG1 concentrations in the same supernatants were not reduced by inhibitor in the cultures (Fig. 6C, D). Although IL-4 can promote B cell proliferation and survival in a PARP14-dependent manner (66), this effect was not likely to be the basis for reduced IgE caused by RBN2759 as the compound did not substantially reduce the cell numbers of these cultures (Table 2).

**Table 2.**
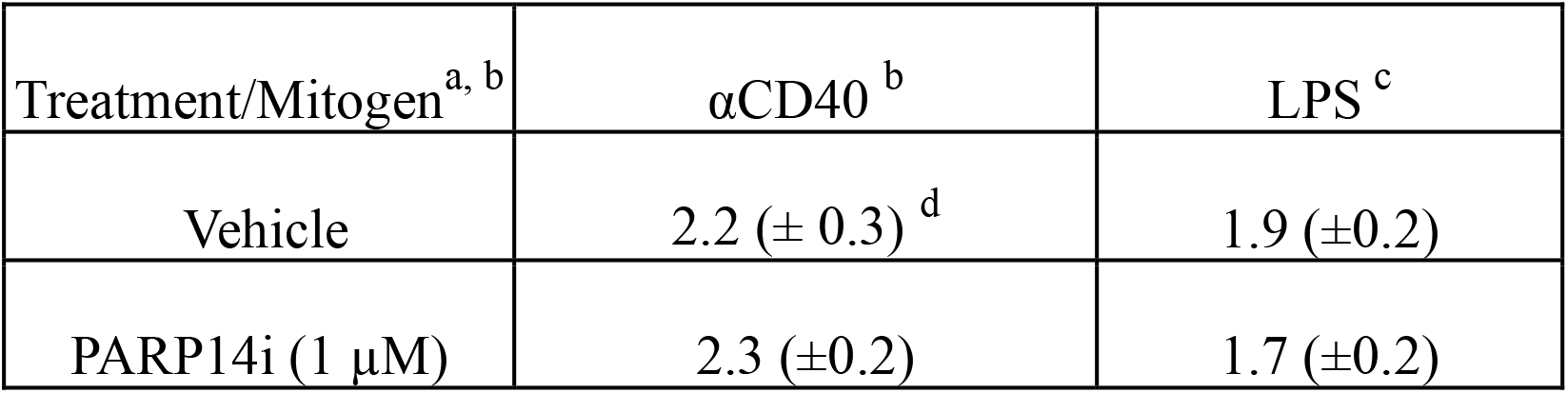
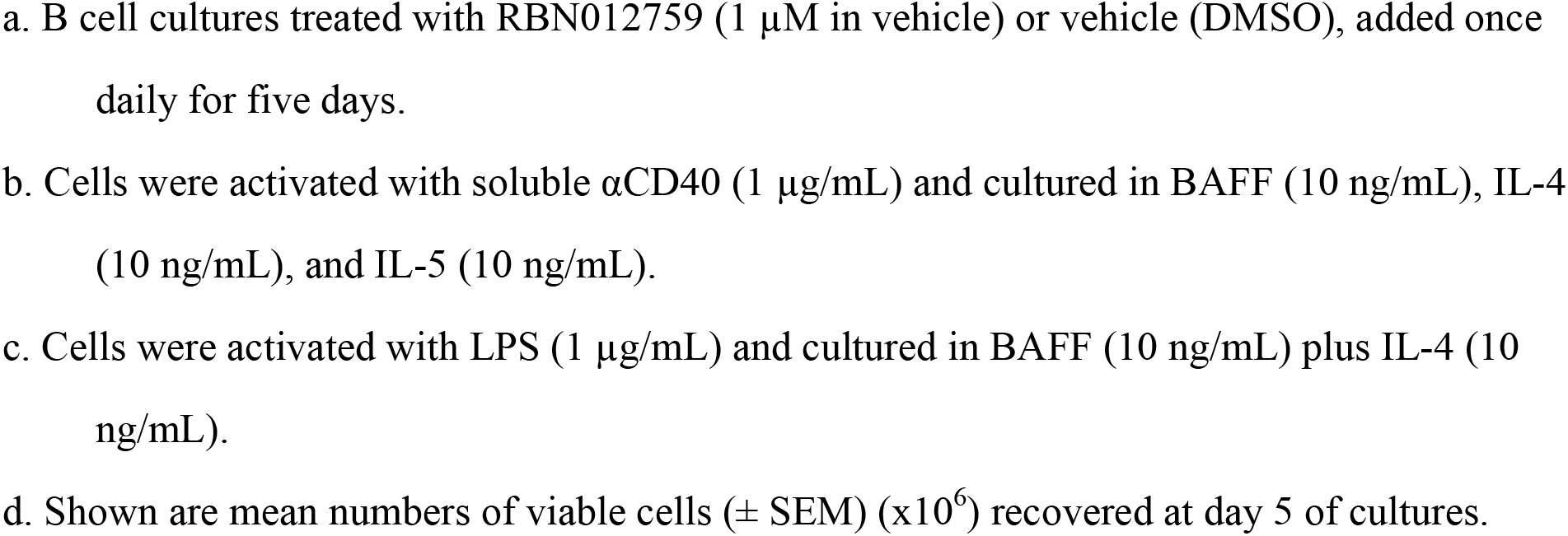

**Fig. 6.**
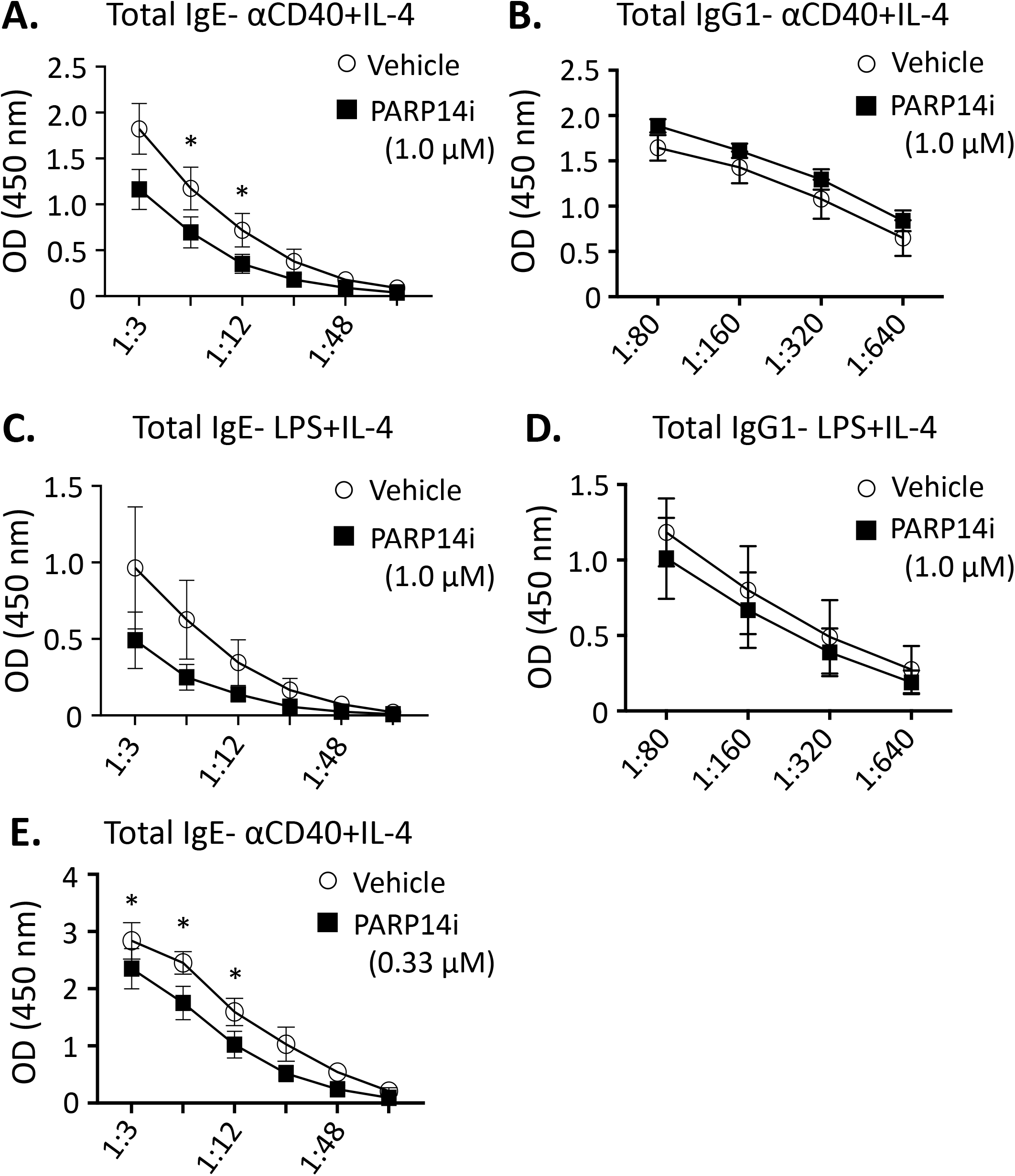
Reciprocal effects of PARP14 inhibition on IgE versus IgG2a production from cultured B cells. (A-E) Measurement of IgE in supernatants from purified B Cell cultures treated with PARP14i or DMSO. (A, B) Measurements of IgE (A) or IgG1 (B) secreted into media 5 d after activation of B cells with αCD40 and culture with IL-4, BAFF, and IL-5. Cultures were treated with 1 μM PARP14i or an equal volume of solvent (DMSO). Shown are means (±SEM) of ELISA values across dilution curves using supernatants of four independent experiments. (C, D) As in (A, B) except purified B cells were activated with LPS (n= 4 independent experiments). A Two-way ANOVA was performed across the curve (p< 0.05). (E) As in (A) except that RBN012759 was used at 0.33 μM.

## DISCUSSION

While a large percentage of human populations has been shown to be reactive to A. *alternata,* patients previously diagnosed with asthma are at higher risk of hospital admission or death upon exposure to *Alternaria* (56, 57). Using an extract of this clinically relevant aeroallergen in a mouse model of allergic lung inflammation, our results show that an inhibitor of catalytic function highly selective for the ADP-ribosyltransferase of PARP14 (alternatively designated ARTD8) is biologically active. Of note, even after having first sensitized subjects by repeated antigen inhalations, treatment with RBN2759 mitigated both the allergen-induced increases of bronchial mucus and IgE. These findings suggest that the approach of inhibiting the catalytic function of PARP14 has therapeutic potential in patients, in that the drug was effective in a recall model in mice that were already sensitized to the aeroallergen.

Prior work had indicated that ovalbumin-induced allergic lung responses were reduced when mice lack PARP14 (36, 37). However, PARP14 has several functional domains other than the conserved catalytic domain towards its C terminus (38, 40, 41, 45, 46). Three modules termed ‘macro domains’-based on similarity to the extension of a large histone variant termed macro-H2a (46, 67, 68) - are situated amino-terminal to the catalytic domain. Crystal structures of PARP14 indicate that its macro domains can bind ADP-ribose [(40, 46). In addition, the amino terminal half of PARP14 contains RNA binding and helicase functions (41). While these domains may contribute to the spectrum of phenotypes observed for PARP14-deficient mice or specific immune cells from them (35, 41), the findings here provide direct support for substantial effects of protein ADP-monoribosylation under physiological conditions in recall allergic lung inflammation.

Treatment of ovalbumin-sensitized *Parp14* −/− mice with the NAD^+^ analogue PJ34 further reduced a subset of cytokine readouts but did not affect some other cytokine levels (37). The published interpretation of this result is at best complex and fraught, as the IC50 of PJ34 for PARP14 (1-10 μM) is ~100-fold less favorable than for PARP1 and PARP2 (48). Moreover, inactivation of PARP1 either by genetic or a more specific pharmacological agent blunted ovalbumin-induced allergic lung inflammation and type 2 cytokines (49–53). Of note, assays of RBN2759 indicated that there is a >>1000-fold higher IC50 for any affect on PARP1 or any other polyPARP, and >300-fold selectivity relative to the other enzymes that catalyze ADPr modifications (55). Although the data show that peak blood concentrations of the agent in this study (~1.3 – 4 μM) might transiently exceed the IC50 of some other PARPs, the trough concentrations were well below the IC50 of all the ADP-ribosyltransferases other than PARP14. Together with the lack of any heterozygote phenotype (Cho SH, Boothby, M, unpublished observations), it is very unlikely that the in vivo target of RBN2759 is either PARP1 or any of the other (ADP-ribosyl)polymerases.

The issue of which ART and PARP enzymes are subject to inhibition of catalysis bears on the risk-benefit balance therapeutic potential of a given agent in the context of specific disease targets. PARP1/2 inhibitors are approved for indications in human cancer but not without dose-limiting toxicities (69–71), for which the duration of treatment will be shorter and the natural history of untreated disease more dire than in allergic diseases or asthma. Thus, what likely are on-target toxicities or side effects do not undermine a favorable risk-benefit relationship in cancer yet would be unsuitable for asthma. The need for sufficient specificity even among mono-ARTs (also termed mono-PARPs) is underscored by work in which a PARP7-specific inhibitor enhances type I interferon responses for potential use in cancer (72, 73). Accordingly, even a less selective mono-ART inhibitor would carry risks of enhancing innate inflammatory cytokines in conditions in which type II (IL-4/5/13-driven) inflammation is the main target. Of note, neither PARP14-deficient nor RBN2759-treated mice have exhibited long- or short-term signals of abnormal health. Moreover, although anti-viral IgA was reduced after infection of *Parp14* −/− mice with mouse-adapted human metapneumovirus, a major cause of viral respiratory disease, illness was not worse (36). Together, these points suggest that chronic treatment with the selective catalytic inhibitor may prove well tolerated in humans as well as effective in the setting of type II inflammation. The findings here suggest that highly selective catalytic inhibitors of this type will also be helpful in research outside the field of allergic disease research. For instance, several separable steps in the replication of diverse viruses require their encoded activity of macro domains for binding to ADPr, or removal of mono-ADPr from unknown protein targets, or both (74–86). Especially since SARS-CoV are among species that use these mechanisms, there will be value in identifying which specific ART enzymes are essential for the endogenous modifications that the virus needs to erase in order to replicate.

### Limitations and unanswered questions

The present body of work both raises intriguing new questions and has inherent limitations. Among these are mechanistic issues, most of all in regards to a sexual disparity (87). In favor of likely efficacy, positive data were obtained for the primary endpoints – mucus scoring and IgE – with both male and female mice and the effect magnitudes for these findings were similar in the two sexes. An encouraging point in support of the findings here is that reductions in BAL and lung inflammation as well as type II cytokines have now been observed in fully independent preliminary replication work with different experimentalists, source of BALB/c male mice (Charles River rather than JAX), mouse colony, and batch of *Alternaria* extract. Nonetheless, later studies would be needed to test whether or not therapeutic efficacy will extend to inbred strains of mice with different patterns of immunity (e.g., C57BL/6; A/J), aeroallergens other than *Alternaria*, or to IgE-mediated or allergic disease processes of skin (e.g., eczema) or gut (e.g., food allergy). In addition, the primary endpoint effects in female mice were not accompanied by decreased cytokine release after ex vivo (re)activation of lung cell suspensions with T cell-specific or non-specific stimuli. This finding leaves open questions as to whether or not the mechanisms are the same in males and females, and what are the cellular and molecular targets in which PARP14 promotes allergic processes. With regards to cellular targets, in vitro data presented here and elsewhere suggest that apart from cytokine production by T cells, PARP14 inhibition changes the pattern of response for both B cells and macrophages (36, 88). However, while the risk of a type 2 statistical error is present, we did not observe a major difference between treatment and control groups with respect to the frequencies of MHC-II^+^ or CD206^+^ macrophages (indices of M1 vs M2) in these experiments on allergic lung inflammation (not shown). As to molecular targets, a fundamental technical barrier has been the difficulty identifying and mapping mono-ADPr modifications on endogenous proteins in physiological settings with full-length PARP14. Thus, cell-free in vitro approaches have provided a plethora of potential candidates (89; Lim J, Cho SH, Boothby M, unpublished observations), but few of these have been validated. Some evidence favors a model in which PARP14-mediated ADP-ribosylation of STAT1 changes its transcriptional effects on monocyte/macrophage-lineage cells (88). In vitro and in vivo, however, major effects of RBN2759 on classical M1-like (M-IFNγ) macrophages may be indirect and due more to a inhibition of the M2-like (M-IL4) population. Accordingly, despite unimpressive results with surface staining for MHC-II and CD206, it is possible that PARP14 inhibition changes the products of alternatively activated macrophages that are key intermediaries in allergic lung inflammation (90). Unfortunately, this avenue of research is so unlikely to be fundable via the US NIH that the probability of developing answers to the questions posed above by that route is vanishingly low or nil despite the promise.

## ACKNOWLEDGEMENTS

We gratefully acknowledge support of the American Asthma Foundation (AAF, formerly Sandler Program in Asthma Research) for the Extension Grant that made possible these studies of the therapeutic potential, as well as that of the Pathology, Microbiology & Immunology Department of VUMC to allow a wrap-up after the end of AAF funds. We also are grateful to Prof. L. Wu and the department for flow cytometry capabilities, the cores of VUMC and Vanderbilt University that were used in this work (Flow Cytometry, Translational Pathology, and Molecular Biology reagents), to S. Toki for helpful technical discussions pertaining to the use of *Alternaria* extract, and D. Nichols to A. M. E. in histologic techniques.

## ABBREVIATIONS USED

ALI: allergic lung inflammation
Ig: immunoglobulin
STAT: Signal Transducer and Activator of Transcription
Th: T helper
IL-: interleukin-
GLT: germline transcript
PARP: Poly-(ADP-ribose) Polymerase
ART: ADP-ribosyltransferase
ADPr: ADP-ribose
CD3, 28, 40, etc: Cluster of Differentiation (molecule #3, 28, 40, etc)
ELISA: enzyme-linked immunosorbent assay
PBS: phosphate-buffered saline
IC (50): inhibitory concentration achieving 50% effect
i.n.: intranasal
Ab: antibody
H+L: heavy and light chains (of Ab molecule)
WT: wild-type
Ag: antigen
MAR: mono ADPr
PAR: poly ADPr
PAS: Periodic Acid Schiff
BAL: broncho-alveolar lavage
TNF: tumor necrosis factor
LPS: lipopolysaccharide
MHC: major histocompatibility complex

## AUTHOR CONTRIBUTIONS and potential conflicts of interest

A. M. E. initiated drafting of this manuscript and, with guidance from S. H. C. and A. L. R., performed experiments and statistical analyses; with guidance from S. H. C. and A. L. R., K. C. performed analyses of samples; K. K. handled analyses of drug concentrations and guidance in dosing and delivery; K. G. analyzed BAL differential counts; L. S. and K. S. generated compound and provided guidance in its use; J. M. and M. N. assisted in design and interpretation of the study as well as providing information on biological data with RBN012759 and editorial input on the manuscript; H. K. coordinated liason among Ribon employees and the experimental team at VUMC, along with input on the manuscript; J. M., M. N., and H. K. shared initial follow-up findings with independent confirmation of the conclusions herein; A. L. R. provided guidance to A. M. E. and K. C. on experiments and their analyses, and input into the manuscript; R. S. P. provided assistance and guidance on the design and interpretation of the study; M. R. B. provided overall guidance and framework, liason with Ribon, wrote the manuscript, and made all final decisions on text and interpretation; S. H. C. provided overall guidance to A. M. E. and K. C., performed experiments, processed and interpreted data, and participated in preparation of the manuscript. K. K., L. S., K. S., J. M., M. N., and H. K. are full-time employees of and hold equity interests in Ribon Therapeutics but were recused from analyses and interpretation of the data and from manuscript preparation; A. M. E., K. C., K. G., A. L. R., R. S. P., and S. H. C. report no potential conflicts of interest; M. R. B. holds equity in Regeneron, Inc., which markets a biologic agent used in treatment of allergic diseases and asthma.

## Notes

### Competing Interest Statement

All authors who are or were employees of Ribon Therapeutics, Inc. would benefit financially if the pharmaceutical agent that is central to this report, or derivatives thereof, were to make it to be an approved therapeutic. Thus, K. K., L. S., K. S., J. M., M. N., and H. K. are full-time employees of and hold equity interests in Ribon Therapeutics but were recused from analyses and interpretation of the data and from manuscript preparation; A. M. E., K. C., K. G., A. L. R., R. S. P., and S. H. C. report no potential conflicts of interest; M. R. B. holds equity in Regeneron, Inc., which markets a biologic agent used in treatment of allergic diseases and asthma.

## References

1. Hekking PP, Wener RR, Amelink M, Zwinderman AH, Bouvy ML, Bel EH. 2015. The prevalence of severe refractory asthma. J Allergy Clin Immunol. 2135:896–902.

2. Loftus PA, Wise SK. 2016. Epidemiology of asthma. Curr Opin Otolaryngol Head Neck Surg. 24:245–249.

3. CDC.gov. 2018. CDC - Asthma - Data and Surveillance - Asthma Surveillance Data. [online] Available at: http://www.cdc.gov/asthma/asthmadata.htm

4. National Center for Health Statistics. Survey Description, National Health Interview Survey, 2017. Hyattsville, Maryland. 2018.

5. Barnes PJ. 2004. Corticosteroid resistance in airway disease. Proc Am Thoracic Soc. 1:264–268.

6. Chung KF. 2013. New treatments for severe treatment-resistant asthma: targeting the right patient. Lancet Respir Med. 1:639–652.

7. Barnes PJ. 2013. Corticosteroid resistance in patients with asthma and chronic obstructive pulmonary disease. J Allergy Clin Immunol. 131:636–645.

8. Lindsay JT, Heaney LG. 2013. Non-adherence in difficult asthma and advances in detection. Expert Rev Respir Med. 2013 Dec; 7(6):607–614.

9. Bender BG. 2016. Nonadherence to asthma treatment: Getting unstuck. J Allerg Clin Immunol: In practice. 4:849–851.

10. Rhodes HL, Thomas P, Sporik R, Holgate ST, Cogswell JJ. 2002. A birth cohort study of subjects at risk of atopy: twenty-two-year follow-up of wheeze and atopic status. Am J Respir Crit Care Med. 165:176–180.

11. Subrata LS, Bizzintino J, Mamessier E, Bosco A, McKenna KL, Wikström ME, Goldblatt J, Sly PD, Hales BJ, Thomas WR, Laing IA, LeSouëf PN, Holt PG. 2009. Interactions between innate antiviral and atopic immunoinflammatory pathways precipitate and sustain asthma exacerbations in children. J Immunol. 183:2793–2800.

12. Holt PG, Sly PD. 2012. Viral infections and atopy in asthma pathogenesis: new rationales for asthma prevention and treatment. Nat Med. 18:726–735.

13. Martinez, Fernando D. 2019. The state of asthma research: considerable advances, but still a long way to go. Am J Respir Crit Care Med. 199:397–399.

14. Wills-Karp M. 1999. Immunologic basis of antigen-induced airway hyperresponsiveness. Annu Rev Immunol. 17:255–281.

15. Finkelman FD, Hogan SP, Hershey GK, Rothenberg ME, Wills-Karp M. 2010. Importance of cytokines in murine allergic airway disease and human asthma. J Immunol. 84:1663–1674.

16. Boonpiyathad T, Sözener ZC, Satitsuksanoa P, Akdis CA. 2019. Immunologic mechanisms in asthma. Semin Immunol. 46:101333.

17. Chesné J, Braza F, Mahay G, Brouard S, Aronica M, Magnan A. 2014. IL-17 in severe asthma. Where do we stand? Am J Respir Crit Care Med. 190:1094–1101.

18. Newcomb DC, Peebles RS Jr. 2013. Th17-mediated inflammation in asthma. Curr Opin Immunol. 25:755–760.

19. Ray A, Kolls JK. 2017. Neutrophilic inflammation in asthma and association with disease severity. Trends Immunol. 38:942–954.

20. Seys SF, Lokwani R, Simpson JL, Bullens DMA. 2019. New insights in neutrophilic asthma. Curr Opin Pulm Med. 25:113–120.

21. Ray A, Camiolo M, Fitzpatrick A, Gauthier M, Wenzel SE. 2020. Are we meeting the promise of endotypes and precision medicine in asthma? Physiol Rev. 100:983–1017.

22. Hamilton D, Lehman H. 2020. Asthma phenotypes as a guide for current and future biologic therapies. Clin Rev Allergy Immunol. 59:160–174.

23. Kühn R, Rajewsky K, Müller W. 1991. Generation and analysis of interleukin-4 deficient mice. Science 254:707–710.

24. Shimoda K, van Deursen J, Sangster MY, Sarawar SR, Carson RT, Tripp RA, Chu C, Quelle FW, Nosaka T, Vignali DA, Doherty PC, Grosveld G, Paul WE, Ihle JN. 1996. Lack of IL-4-induced Th2 response and IgE class switching in mice with disrupted Stat6 gene. Nature 380:630–633.

25. Mochizuki M, Bartels J, Mallet AI, Christophers E, Schröder JM. 1998. IL-4 induces eotaxin: a possible mechanism of selective eosinophil recruitment in helminth infection and atopy. J Immunol. 160:60–68.

26. Cohn L, Homer RJ, MacLeod H, Mohrs M, Brombacher F, Bottomly K. 1999. Th2-induced airway mucus production is dependent on IL-4Ralpha, but not on eosinophils. J Immunol. 162:6178–6183.

27. Teran LM, Mochizuki M, Bartels J, Valencia EL, Nakajima T, Hirai K, Schröder JM. 1999. Th1- and Th2-type cytokines regulate the expression and production of eotaxin and RANTES by human lung fibroblasts. Am J Respir Cell Mol Biol. 20:777–786.

28. Mathew A, MacLean JA, DeHaan E, Tager AM, Green FH, Luster AD. 2001. Signal transducer and activator of transcription 6 controls chemokine production and T helper cell type 2 cell trafficking in allergic pulmonary inflammation. J Exp Med. 193:1087–1096.

29. Tomkinson A, Duez C, Cieslewicz G, Pratt JC, Joetham A, Shanafelt MC, Gundel R, Gelfand EW. 2001. A murine IL-4 receptor antagonist that inhibits IL-4- and IL-13-induced responses prevents antigen-induced airway eosinophilia and airway hyperresponsiveness. J Immunol. 166:5792–5800.

30. Hoshino A, Tsuji T, Matsuzaki J, Jinushi T, Ashino S, Teramura T, Chamoto K, Tanaka Y, Asakura Y, Sakurai T, Mita Y, Takaoka A, Nakaike S, Takeshima T, Ikeda H, Nishimura T. 2004. STAT6-mediated signaling in Th2-dependent allergic asthma: critical role for the development of eosinophilia, airway hyper-responsiveness and mucus hypersecretion, distinct from its role in Th2 differentiation. Int Immunol. 16:1497–1505.

31. Snapper CM, Paul WE. 1987. Interferon-gamma and B cell stimulatory factor-1 reciprocally regulate Ig isotype production. Science 236:944–947.

32. Rothman P, Li S, Gorham B, Glimcher L, Alt F, Boothby M. 1991. Identification of a conserved lipopolysaccharide-plus-interleukin-4-responsive element located at the promoter of germ line epsilon transcripts. Mol Cell Biol. 11:5551–5561.

33. Linehan LA, Warren WD, Thompson PA, Grusby MJ, Berton MT. 1998. STAT6 is required for IL-4-induced germline Ig gene transcription and switch recombination. J Immunol. 161:302–310.

34. Sarfati M, Wakahara K, Chapuy L, Delespesse G. 2015. Mutual Interaction of Basophils and T Cells in Chronic Inflammatory Diseases. Front Immunol. 6:399.

35. Goenka S, Boothby M. 2006. Selective potentiation of Stat-dependent gene expression by collaborator of Stat6 (CoaSt6), a transcriptional cofactor. Proc Natl Acad Sci USA 103:4210–4215.

36. Cho SH, Raybuck A, Wei M, Erickson J, Nam KT, Cox RG, Trochtenberg A, Thomas JW, Williams J, Boothby M. 2013. B cell-intrinsic and -extrinsic regulation of antibody responses by PARP14, an intracellular (ADP-ribosyl)transferase. J Immunol. 191:3169–3178.

37. Mehrotra P, Hollenbeck A, Riley JP, Li F, Patel RJ, Akhtar N, Goenka S. 2013. Poly (ADP-ribose) polymerase 14 and its enzyme activity regulates T(H)2 differentiation and allergic airway disease. J Allergy Clin Immunol. 131:521–531.

38. Goenka S, Cho SH, Boothby M. 2007. Collaborator of Stat6 (CoaSt6)-associated poly(ADP-ribose) polymerase activity modulates Stat6-dependent gene transcription. J Biol Chem. 282:18732–19739.

39. Kleine H, Poreba E, Lesniewicz K, Hassa PO, Hottiger MO, Litchfield DW, Shilton BH, Lüscher B. 2008. Substrate-assisted catalysis by PARP10 limits its activity to mono-ADP-ribosylation. Mol Cell. 32:57–69.

40. Forst AH, Karlberg T, Herzog N, Thorsell AG, Gross A, Feijs KL, Verheugd P, Kursula P, Nijmeijer B, Kremmer E, Kleine H, Ladurner AG, Schüler H, Lüscher B. 2013. Recognition of mono-ADP-ribosylated ARTD10 substrates by ARTD8 macrodomains. Structure 21:462–475.

41. Iqbal MB, Johns M, Cao J, Liu Y, Yu SC, Hyde GD, Laffan MA, Marchese FP, Cho SH, Clark AR, Gavins FN, Woollard KJ, Blackshear PJ, Mackman N, Dean JL, Boothby M, Haskard DO. 2014. PARP-14 combines with tristetraprolin in the selective post-transcriptional control of macrophage tissue factor expression. Blood 124:3646–3655.

42. Ekblad T, Lindgren AE, Andersson CD, Caraballo R, Thorsell AG, Karlberg T, Spjut S, Linusson A, Schüler H, Elofsson M. 2015. Towards small molecule inhibitors of mono-ADP-ribosyltransferases. Eur J Med Chem. 95:546–551.

43. Thorsell AG, Ekblad T, Karlberg T, Löw M, Pinto AF, Trésaugues L, Moche M, Cohen MS, Schüler H. 2017. Structural basis for potency and promiscuity in Poly(ADP-ribose) Polymerase (PARP) and Tankyrase inhibitors. J Med Chem. 60:1262–1271.

44. Holechek J, Lease R, Thorsell AG, Karlberg T, McCadden C, Grant R, Keen A, Callahan E, Schüler H, Ferraris D. 2018. Design, synthesis and evaluation of potent and selective inhibitors of mono-(ADP-ribosyl)transferases PARP10 and PARP14. Bioorg Med Chem Lett. 28:2050–2054.

45. Gupte R, Liu Z, Kraus WL. 2017. PARPs and ADP-ribosylation: recent advances linking molecular functions to biological outcomes. Genes Dev. 31:101–126.

46. Schreiber V, Dantzer F, Ame JC, de Murcia G. 2006. Poly(ADP-ribose): novel functions for an old molecule. Nat Rev Mol Cell Biol. 7:517–528.

47. Di Girolamo M, Fabrizio G. 2019. Overview of the mammalian ADP-ribosyl-transferases clostridia toxin-like (ARTCs) family. Biochem Pharmacol. 167:86–96.

48. Wigle TJ, Church WD, Majer CR, Swinger KK, Aybar D, Schenkel LB, Vasbinder MM, Brendes A, Beck C, Prahm M, Wegener D, Chang P, Kuntz KW. 2020. Forced self-modification assays as a strategy to screen monoPARP enzymes. SLAS Discov. 25:241–252.

49. Boulares AH, Zoltoski AJ, Sherif ZA, Jolly P, Massaro D, Smulson ME. 2003. Gene knockout or pharmacological inhibition of poly(ADP-ribose) polymerase-1 prevents lung inflammation in a murine model of asthma. Am J Respir Cell Mol Biol. 28:322–329.

50. Oumouna M, Datta R, Oumouna-Benachour K, Suzuki Y, Hans C, Matthews K, Fallon K, Boulares H. 2006. Poly(ADP-ribose) polymerase-1 inhibition prevents eosinophil recruitment by modulating Th2 cytokines in a murine model of allergic airway inflammation: a potential specific effect on IL-5. J Immunol. 177:6489–6496.

51. Ghonim MA, Pyakurel K, Ibba SV, Al-Khami AA, Wang J, Rodriguez P, Rady HF, El-Bahrawy AH, Lammi MR, Mansy MS, Al-Ghareeb K, Ramsay A, Ochoa A, Naura AS, Boulares AH. 2015. PARP inhibition by olaparib or gene knockout blocks asthma-like manifestation in mice by modulating CD4(+) T cell function. J Transl Med. 13:225.

52. Ghonim MA, Pyakurel K, Ibba SV, Wang J, Rodriguez P, Al-Khami AA, Lammi MR, Kim H, Zea AH, Davis C, Okpechi S, Wyczechowska D, Al-Ghareeb K, Mansy MS, Ochoa A, Naura AS, Boulares AH. 2015. PARP is activated in human asthma and its inhibition by olaparib blocks house dust mite-induced disease in mice. Clin Sci (Lond). 129:951–962.

53. Sethi GS, Sharma S, Naura AS. 2019. PARP inhibition by olaparib alleviates chronic asthma-associated remodeling features via modulating inflammasome signaling in mice. IUBMB Life 71:1003–1013.

54. Wigle TJ, Blackwell DJ, Schenkel LB, Ren Y, Church WD, Desai HJ, Swinger KK, Santospago AG, Majer CR, Lu AZ, Niepel M, Perl NR, Vasbinder MM, Keilhack H, Kuntz KW. 2020. In vitro and cellular probes to study PARP enzyme target engagement. Cell Chem Biol. 27:877–887.

55. Schenkel L, Molina J, Swinger K, Abo R, Blackwell D, Cheung A, Church W, Kuplast-Barr K, Lu A, Minissale E, Niepel M, Vasbinder M, Wigle T, Richon V, Keilhack H, Kuntz KW. 2020. A potent and selective PARP14 inhibitor decreases pro-tumor macrophage function and elicits inflammatory responses in tumor explants. Cancer Res. Abstract 1038: DOI: 10.1158/1538-7445.AM2020-1038.

56. Neukirch C, Henry C, Leynaert B, Liard R, Bousquet J, Neukirch F. 1999. Is sensitization to *Alternaria alternata* a risk factor for severe asthma? A population-based study. J Allergy Clin Immunol 103:709–711.

57. Bush RK, Prochnau JJ. 2004. *Alternaria*-induced asthma. J Allergy Clin Immunol 113:227–?.

58. Newcomb DC, Boswell MG, Reiss S, Zhou W, Goleniewska K, Toki S, Harintho MT, Lukacs NW, Kolls JK, Peebles RS Jr. 2013. IL-17A inhibits airway reactivity induced by respiratory syncytial virus infection during allergic airway inflammation. Thorax 68:717–723.

59. Lee K, Gudapati P, Dragovic S, Spencer C, Joyce S, Killeen N, Magnuson MA, Boothby M. 2010. Mammalian target of rapamycin protein complex 2 regulates differentiation of Th1 and Th2 cell subsets via distinct signaling pathways. Immunity 32:743–753.

60. Dunican EM, Elicker BM, Gierada DS, Nagle SK, Schiebler ML, Newell JD, Raymond WW, Lachowicz-Scroggins ME, Di Maio S, Hoffman EA, Castro M, Fain SB, Jarjour NN, Israel E, Levy BD, Erzurum SC, Wenzel SE, Meyers DA, Bleecker ER, Phillips BR, Mauger DT, Gordon ED, Woodruff PG, Peters MC, Fahy JV; National Heart Lung and Blood Institute (NHLBI) Severe Asthma Research Program (SARP). 2018. Mucus plugs in patients with asthma linked to eosinophilia and airflow obstruction. J Clin Invest. 128:997–1009.

61. Dunican EM, Watchorn DC, Fahy JV. 2018. Autopsy and imaging studies of mucus in asthma. Lessons learned about disease mechanisms and the role of mucus in airflow obstruction. Ann Am Thorac Soc. 15(Suppl 3): S184–S191.

62. Humbles AA, Lloyd CM, McMillan SJ, Friend DS, Xanthou G, McKenna EE, Ghiran S, Gerard NP, Yu C, Orkin SH, Gerard C. 2004. A critical role for eosinophils in allergic airways remodeling. Science 305:1776–1779.

63. Aceves SS, Broide DH. 2008. Airway fibrosis and angiogenesis due to eosinophil trafficking in chronic asthma. Curr Mol Med. 8:350–358.

64. Foster PS, Maltby S, Rosenberg HF, Tay HL, Hogan SP, Collison AM, Yang M, Kaiko GE, Hansbro PM, Kumar RK, Mattes J. 2017. Modeling T_H_ 2 responses and airway inflammation to understand fundamental mechanisms regulating the pathogenesis of asthma. Immunol Rev. 278:20–40.

65. Mizutani N, Nabe T, Yoshino S. 2014. IL-17A promotes the exacerbation of IL-33-induced airway hyperresponsiveness by enhancing neutrophilic inflammation via CXCR2 signaling in mice. J Immunol. 192:1372–1384.

66. Cho SH, Goenka S, Henttinen T, Gudapati P, Reinikainen A, Eischen CM, Lahesmaa R, Boothby M. 2009. PARP-14, a member of the B aggressive lymphoma family, transduces survival signals in primary B cells. Blood 113:2416–2425.

67. Karras GI, Kustatscher G, Buhecha HR, Allen MD, Pugieux C, Sait F, Bycroft M, Ladurner AG. 2005. The macro domain is an ADP-ribose binding module. EMBO J. 24:1911–1920.

68. Feijs KL, Forst AH, Verheugd P, Lüscher B. 2013. Macrodomain-containing proteins: regulating new intracellular functions of mono(ADP-ribosyl)ation. Nat Rev Mol Cell Biol. 14:443–451.

69. Chan CY, Tan KV, Cornelissen B. 2020. PARP inhibitors in cancer diagnosis and therapy. Clin Cancer Res. 2020 Oct 20:clincanres.2766.2020. doi: 10.1158/1078-0432.CCR-2-2766.

70. Curtin NJ, Szabo C. 2020. Poly(ADP-ribose) polymerase inhibition: past, present and future. Nat Rev Drug Discov. 19:711–736.

71. Eakin CM, Ewongwo A, Pendleton L, Monk BJ, Chase DM. 2020. Real world experience of poly (ADP-ribose) polymerase inhibitor use in a community oncology practice. Gynecol Oncol. 159:112–117.

72. Yamada T, Horimoto H, Kameyama T, Hayakawa S, Yamato H, Dazai M, Takada A, Kida H, Bott D, Zhou AC, Hutin D, Watts TH, Asaka M, Matthews J, Takaoka A. 2016. Constitutive aryl hydrocarbon receptor signaling constrains type I interferon-mediated antiviral innate defense. Nat Immunol. 17:687–694.

73. https://pipelinereview.com/index.php/2019090572150/Small-Molecules/Ribon-Therapeutics-Announces-Dosing-of-First-Patient-in-Phase-1-Clinical-Trial-of-RBN-2397-a-First-In-Class-PARP7-Inhibitor-Ribon-Therapeutics-Announces-Dosing-of-First-P.html

74. Dunstan MS, Barkauskaite E, Lafite P, Knezevic CE, Brassington A, Ahel M, Hergenrother PJ, Leys D, Ahel I. 2012. Structure and mechanism of a canonical poly(ADP-ribose) glycohydrolase. Nat Commun. 3:878.

75. Egloff MP, Malet H, Putics A, Heinonen M, Dutartre H, Frangeul A, Gruez A, Campanacci V, Cambillau C, Ziebuhr J, Ahola T, Canard B. 2006. Structural and functional basis for ADP-ribose and poly(ADP-ribose) binding by viral macro domains. J Virol. 80:8493–8502.

76. Eriksson KK, Cervantes-Barragán L, Ludewig B, Thiel V. 2008. Mouse hepatitis virus liver pathology is dependent on ADP-ribose-1’’-phosphatase, a viral function conserved in the alpha-like supergroup. J Virol. 82:12325–12334.

77. Neuvonen M, Ahola T. 2009. Differential activities of cellular and viral macro domain proteins in binding of ADP-ribose metabolites. J Mol Biol. 385:212–225.

78. Jankevicius G, Hassler M, Golia B, Rybin V, Zacharias M, Timinszky G, Ladurner AG. 2013. A family of macrodomain proteins reverses cellular mono-ADP-ribosylation. Nat Struct Mol Biol. 20:508–14.

79. Rosenthal F, Feijs KL, Frugier E, Bonalli M, Forst AH, Imhof R, Winkler HC, Fischer D, Caflisch A, Hassa PO, Lüscher B, Hottiger MO. 2013. Macrodomain-containing proteins are new mono-ADP-ribosylhydrolases. Nat Struct Mol Biol. 20:502–507.

80. Daugherty MD, Young JM, Kerns JA, Malik HS. 2014. Rapid evolution of PARP genes suggests a broad role for ADP-ribosylation in host-virus conflicts. PLoS Genet. 10:e1004403.

81. Atasheva S, Frolova EI, Frolov I. 2014.Interferon-stimulated poly(ADP-Ribose) polymerases are potent inhibitors of cellular translation and virus replication. J Virol. 88:2116–2130.

82. Cho CC, Lin MH, Chuang CY, Hsu CH. 2016. Macro domain from Middle Eastern Respiratory Syndrome Coronavirus (MERS-CoV) is an efficient ADP-ribose binding module: Crystal structure and biochemical studies. J Biol Chem. 291:4894–4902.

83. Li C, Debing Y, Jankevicius G, Neyts J, Ahel I, Coutard B, Canard B. 2016. Viral Macro domains reverse protein ADP-Ribosylation. J Virol. 90:8478–8486.

84. McPherson RL, Abraham R, Sreekumar E, Ong SE, Cheng SJ, Baxter VK, Kistemaker HA, Filippov DV, Griffin DE, Leung AK. 2017. ADP-ribosylhydrolase activity of Chikungunya virus macrodomain is critical for virus replication and virulence. Proc Natl Acad Sci USA 114:1666–1671.

85. Abraham R, Hauer D, McPherson RL, Utt A, Kirby IT, Cohen MS, Merits A, Leung AKL, Griffin DE. 2018. ADP-ribosyl-binding and hydrolase activities of the alphavirus nsP3 macrodomain are critical for initiation of virus replication. Proc Natl Acad Sci USA 115:E10457–E10466.

86. Grunewald ME, Chen Y, Kuny C, Maejima T, Lease R, Ferraris D, Aikawa M, Sullivan CS, Perlman S, Fehr AR. 2019. The coronavirus macrodomain is required to prevent PARP-mediated inhibition of virus replication and enhancement of IFN expression. PLoS Pathog. 15:e1007756.

87. Cephus JY, Stier MT, Fuseini H, Yung JA, Toki S, Bloodworth MH, Zhou W, Goleniewska K, Zhang J, Garon SL, Hamilton RG, Poloshukin VV, Boyd KL, Peebles RS Jr, Newcomb DC. 2017. Testosterone attenuates group 2 innate lymphoid cell-mediated airway inflammation. Cell Rep. 21:2487–2499.

88. Iwata H, Goettsch C, Sharma A, Ricchiuto P, Goh WW, Halu A, Yamada I, Yoshida H, Hara T, Wei M, Inoue N, Fukuda D, Mojcher A, Mattson PC, Barabási AL, Boothby M, Aikawa E, Singh SA, and Aikawa M. 2016. PARP9 and PARP14 cross-regulate macrophage activation via STAT1 ADP-ribosylation. Nat Commun. 7:12849.

89. Feijs KL, Kleine H, Braczynski A, Forst AH, Herzog N, Verheugd P, Linzen U, Kremmer E, Lüscher B. 2013. ARTD10 substrate identification on protein microarrays: regulation of GSK3β by mono-ADP-ribosylation. Cell Commun Signal. 11(1):5.

90. Dasgupta P, Keegan AD. 2012. Contribution of alternatively activated macrophages to allergic lung inflammation: a tale of mice and men. J Innate Immun. 4:478–88.

